# Zika Virus Infection at Different Pregnancy Stages: Anatomopathological Findings, Target Cells and Viral Persistence in Placental Tissues

**DOI:** 10.1101/370528

**Authors:** Lucia de Noronha, Camila Zanluca, Marion Burger, Andreia Akemi Suzukawa, Marina Azevedo, Patricia Z. Rebutini, Iolanda Maria Novadzki, Laurina Setsuko Tanabe, Mayra Marinho Presibella, Claudia Nunes Duarte dos Santos

## Abstract

Zika virus (ZIKV) infection in humans has been associated with congenital malformations and other neurological disorders, such as Guillain-Barré syndrome. The mechanism(s) of ZIKV intrauterine transmission, the cell types involved, the most vulnerable period of pregnancy for severe outcomes from infection and other physiopathological aspects remain unknown. In this study, we analyzed placental samples obtained at the time of delivery from a group of twenty-four women diagnosed with ZIKV infection during the first, second or third trimesters of pregnancy. Villous immaturity was the main histological finding in the placental tissues, although placentas without alterations were also frequently observed. Significant enhancement of the number of syncytial sprouts was observed in the placentas of women infected during the third trimester, indicating the development of placental abnormalities after ZIKV infection. Hyperplasia of Hofbauer cells (HCs) was also observed in these third-trimester placental tissues, and remarkably, HCs were the only ZIKV-positive fetal cells found in the placentas studied that persisted until birth, as revealed by immunohistochemical (IHC) analysis. Thirty-three percent of women infected during pregnancy delivered infants with congenital abnormalities, although no pattern correlating the gestational stage at infection, the IHC positivity of HCs in placental tissues and the presence of congenital malformations at birth was observed. Placental tissue analysis enabled us to confirm maternal ZIKV infection in cases where serum from the acute infection phase was not available, which reinforces the importance of this technique in identifying possible causal factors of birth defects. The results we observed in the samples from naturally infected pregnant women may contribute to the understanding of some aspects of the pathophysiology of ZIKV.

## 1 Introduction

Zika virus (ZIKV) is an emergent arthropod-borne virus that belongs to the genus *Flavivirus* of the *Flaviviridae* family (ICTV. International Comittee on Taxonomy of Viruses, 2017). This virus is primarily transmitted through the bite of the *Aedes* mosquito (Zanluca and Duarte dos Santos, 2016). Unlike most other flaviviruses, however, person-to-person ZIKV transmission is possible, although the contribution of this transmission mode to maintaining an epidemic is unclear. Transmission by sexual and perinatal interactions and from blood and platelet transfusions has been described (Miner and Diamond, 2017; Mlakar et al., 2016; Noronha et al., 2016).

In general, ZIKV infection in humans is characterized as a self-limiting disease, and the most frequent signs and symptoms are low fever, myalgia, rash, arthralgia, headache and conjunctival hyperemia (Duffy et al., 2009; Zanluca et al., 2015). Nevertheless, cases of neurological manifestations, such as Guillain-Barré syndrome (Beckham et al., 2016; Noronha et al., 2016; Schuler-Faccini et al., 2016), have been reported in patients diagnosed with ZIKV. In addition, ZIKV infection during pregnancy has been associated with fetal malformations. Brain microcalcification and other central nervous system disorders, ocular abnormalities, and arthrogryposis are all a part of congenital Zika syndrome (Brasil et al., 2016; Melo et al., 2016; Schuler-Faccini et al., 2016). By March 17, 2017, thirty-one countries or territories in the Americas had reported central nervous system malformations that were potentially associated with ZIKV infection, and Brazil is the most affected country to date (WHO. World Health Organization, 2017).

Since June 2015, we have been receiving samples of serum, urine and other body fluids for ZIKV diagnosis. Additionally, during the peak of the ZIKV outbreak in Brazil, in agreement with the local health authorities, most pregnant women in Paraná State suspected of having ZIKV infection were monitored. Samples of tissues, such as the placenta and umbilical cord, as well as fetal tissues (in the case of stillbirths), all of which were collected at the time of delivery, were sent to our laboratory for analysis

Here, we present a case series in which we analyzed placental tissues from women infected with ZIKV at different pregnancy stages, focusing on the anatomopathological and morphometric findings, target cells and viral persistence.

## 2 Material and methods

### 2.1 Patients and ethics approval

This study was approved by Fiocruz and the Brazilian National Ethics Committee of Human Experimentation under the number CAAE: 42481115.7.0000.5248. Since our laboratory is a Reference Center for the Diagnosis of Emerging Viruses of the Brazilian Ministry of Health, we obtained the waiver for the written informed consent to work with these samples. We are in compliance we all the ethical principles and our study was approved by the above mentioned Ethical Committee.

During the ZIKV outbreak in Brazil (2015-2016), 331 pregnant women exhibiting ZIKV infection-compatible symptoms presented to the Public Health Units in Paraná State. Two hundred ninety-two of them tested negative for ZIKV infection, and 39 cases were laboratory-confirmed for ZIKV by RT-PCR. From the positive cases, two patients had spontaneous abortions, and the remaining 37 women were monitored during pregnancy until the time of delivery.

This report describes the anatomopathological and morphometric findings from the placental tissues of 24 pregnant women with confirmed ZIKV infection, i.e., those with positive results by RT-qPCR or immunohistochemical (IHC) analysis of placental tissue.

Fragments of placental tissue were collected immediately after delivery and were frozen. The presence of the ZIKV genome in the samples was investigated by RT-qPCR, and when appropriate, the presence of antibodies in the serum samples was screened with an ELISA, as described below. An additional fragment of the placenta was kept in buffered formalin for further anatomopathological, morphometric and IHC analyses. The clinical condition of the newborns was evaluated immediately after delivery (no follow-up was performed).

### 2.2 RNA extraction and viral genome detection

Viral RNA was isolated from 140 μL of biological fluids (serum, cerebrospinal fluid, lacrimal fluid, saliva or urine) using a QIAamp Viral RNA Mini Kit (Qiagen, Hilden, Germany) according to the manufacturer’s protocol. RNA extraction from unfixed placenta or viscera was performed with an RNeasy Mini Kit (Qiagen), while RNA from formalin-fixed paraffin-embedded (FFPE) tissues was extracted using the ReliaPrep^TM^ FFPE Total RNA Miniprep System (Promega, Madison, USA). RNA was stored at − 80°C until use.

The primers and probes were synthesized and purified by Integrated DNA Technologies (IDT, Coralville, USA). The reporter dye 5-FAM was used for the probes. All real-time assays were performed with the GoTaq Probe 1-Step RT-qPCR System (Promega) with amplification by a LightCycler 96 instrument (Roche, Mannheim, Germany).

The ZIKV genome was detected according to the protocol of Lanciotti et al. (Lanciotti et al., 2008), which used 5 μL of RNA per 20 μL reaction, while the dengue virus (DENV) serotypes were detected according to the protocol of Johnson et al. (Johnson et al., 2005). Human RNAse P was used as an endogenous control (Emery et al., 2004). Positive controls for ZIKV and DENV were obtained by virus isolation from Brazilian patients sera in C6/36 or Vero E6 cells.

### 2.3 Antibody detection

Anti-ZIKV IgM in serum samples was detected with an in-house IgM capture enzyme-linked immunosorbent assay (MAC-ELISA) using β-propiolactone-inactivated ZIKV and MOCK (from noninfected cells) cell cultures as antigens according to the Centers for Disease Control and Prevention (CDC. Centers for Disease Control and Prevention, 2017) guidelines, with minor modifications. A humanized anti-flavivirus monoclonal antibody (MAb) was kindly provided by the CDC and was used as a positive control. Serum samples were also tested for anti-DENV antibodies using either the Panbio Dengue IgM Capture ELISA or the Panbio Dengue IgG Indirect ELISA (Alere, Brisbane, Australia) following the manufacturer’s instructions.

### 2.4 Anatomopathological, immunohistochemical and morphometric analysis

Deparaffinization and rehydration of the FFPE placental samples were performed using xylene and ethanol baths, respectively, and hydrogen peroxide/methanol was used to block endogenous peroxidase activity. Then, the sections were incubated with anti-flavivirus (4G2) or specific anti-ZIKV (produced at ICC/Fiocruz-PR) MAb and then with biotin-free polyvalent HRP (Spring Bioscience, Pleasanton, USA). Finally, the slides were incubated with a freshly prepared substrate mixture (DAB, DakoCytomation) and counterstained with Mayer’s hematoxylin.

The anti-ZIKV MAb did not show any crossreactivity with other flaviviruses (dengue serotypes 1-4 and the yellow fever, Saint Louis encephalitis and West Nile viruses) in a cell culture model (Figure S1). Additionally, 73 placental samples of other etiologies were tested by IHC using anti-ZIKV MAb, and no crossreactivity was observed. Also, RT-qPCR yielded negative results in most of these samples.

Negative controls were obtained either by omitting the primary antibody from the incubation step or by using an unrelated MAb against Chikungunya virus (CHIKV), produced at ICC/Fiocruz-PR, as the primary antibody (Figure S2).

To better identify Hofbauer cells (HCs), the FFPE placental tissue specimens were incubated with the primary antibody anti-human CD163 (Invitrogen, Carlsbad, USA) by using the same protocol.

The placenta slides (three slides from the placental disc, one slide from the chorioamniotic membrane and one slide from umbilical cord) were observed with an Olympus^™^ BX50 optical microscope (Tokyo, Japan), and photomicrographs were acquired in a high-power field (400×) using a Zeiss Axio Scan.Z1^TM^ scanner (Carl Zeiss, Thornwood, USA) for both routine hematoxylin-eosin (H&E) staining and IHC staining. Villus maturation was evaluated by considering the villus morphology and the gestational age at delivery.

Morphometric analyses were performed in placental tissue specimens from mothers who were infected with ZIKV during the third trimester of pregnancy (n = 6). A negative control group (n = 5) was used for comparison (Baurakiades et al., 2011). For each case, 30 high-power fields (HPFs = 400×) were randomly selected, and the villi, CD163+ HCs, knots and sprouts in each field were counted. The mean values of these parameters over the 30 fields were used for the statistical analyses.

Statistical analyses were performed using GraphPad Prism 5.03 software. The Zika-infected and uninfected control groups were compared using the Mann-Whitney test. Differences were considered to be statistically significant at *p*<0.05.

## 3 Results

This report describes the evaluation of the 24 cases of ZIKV infection confirmed in pregnant women at different gestational periods (Table 1).

**Table 1.** Pathological features and laboratory findings from 24 cases of Zika virus infection confirmed during pregnancy

| TIME OF INFECTION<br>- Gestational Trimester<br>/ Probable month of<br>infection* |  | PATIENT<br>CODE | RT-PCR<br>(serum) | RT-PCR<br>(urine) | ANTI-ZIKV IgM<br>(serum) | ANTI-DENV<br>Serology | IHC<br>Placenta<br>(FFPE) | RT-PCR<br>Placenta<br>(FFPE) | RT-PCR<br>Placenta<br>(unfixed) | GESTATIONAL AGE /<br>PATHOLOGICAL<br>FEATURES | CONGENITAL<br>DISORDER | OUTCOME/<br>DELIVERY<br>GESTATIONAL<br>AGE | NEWBORN SAMPLES -<br>Additional observations |
| --- | --- | --- | --- | --- | --- | --- | --- | --- | --- | --- | --- | --- | --- |
| FIRST<br>TRIMESTER<br>(weeks 1-13) | 2<sup>nd</sup> | LRV/16 1350 | + | - | I | IgM-/IgG- | - | NT | - | Third trimester / umbilical<br>artery agenesis | YES – microcephaly,<br>ventriculomegaly,<br>calcifications | Newborn/preterm (36<br>w) | CSF, urine, lacrimal fluid<br>and saliva RT-PCR- |
|  | 3<sup>rd</sup> | LRV/16 1318 | + | - | + | IgM-/IgG+ | - | NT | - | Third trimester / without<br>pathological changes | NO | Newborn/term | CSF, Lacrimal fluid and<br>saliva RT-PCR-<br>Serum RT-PCR-/IgM- |
|  | 1<sup>st</sup> | LRV/15 578 | NA | NA | NA | NA | - † | + | NA | Third trimester / villous<br>immaturity | YES – microcephaly,<br>arthrogryposis, perinatal<br>death | Newborn/term | Brain tissue RT-PCR+ |
|  | 1<sup>st</sup> | LRV/16 1317 | + | NA | + | IgM-/IgG+ | - | NT | - | Third trimester / without<br>pathological changes | NO | Newborn/term | NA |
|  | 2<sup>nd</sup> | LRV/15 572 | NA | NA | NA | NA | + | + | NA | First trimester / chronic<br>villitis TORCH-like | NO | Spontaneous abortion/<br>12 w | NA |
| SECOND<br>TRIMESTER<br>(weeks 14-26) | 6<sup>th</sup> | LRV/16 870 | + | NA | + | IgM-/IgG+ | - | - | - | Third trimester / without<br>pathological changes | NO | Newborn/term | NA |
|  | 4<sup>th</sup> | LRV/16 1065 | + | - | + | IgM-/IgG+ | - | NT | - | Third trimester / without<br>pathological changes | NO | Newborn/term | Lacrimal fluid and saliva<br>RT-PCR-.<br>Serum RT-PCR-/IgM+<br>Colostrum IgM+ |
|  | 5<sup>th</sup> | LRV/16 933 | + | - | I | IgM-/IgG+ | - | - | + | Third trimester / without<br>pathological changes | NO | Newborn/term | NA |
|  | 6<sup>th</sup> | LRV/16 1068 | + | - | + | IgM-/IgG+ | - | NT | + | Third trimester / mild acute<br>funisitis | NO | Newborn/term | Lacrimal fluid and saliva<br>RT-PCR- |
|  | 6<sup>th</sup> | LRV/16 995 | + | NA | I | IgM-/IgG- | - | - | - | Third trimester / without<br>pathological changes | YES – Spina bifida | Newborn/term | NA |
|  | 5<sup>th</sup> | LRV/16 1029 | + | - | + | IgM-/IgG+ | - | - | - | Third trimester / without<br>pathological changes | NO | Newborn/term | Lacrimal fluid, saliva,<br>serum RT-PCR-/IgM-<br>Colostrum IgM+ |
|  | 5<sup>th</sup> | LRV/16 1004 | + | - | + | IgM-/IgG+ | - | - | + | Third trimester / without<br>pathological changes | NO | Newborn/term (40 w) | Serum RT-PCR-/IgM- |
|  | 6<sup>th</sup> | LRV/16 845 | + | NA | - | IgM-/IgG- | + | - | - | Third trimester / villous<br>immaturity | NO | Newborn/term | NA |
| THIRD<br>TRIMESTER<br>(weeks 27-40) | 8<sup>th</sup> | LRV/16 927 | NA | - | NA | NA | + | - | - | Third trimester / without<br>pathological changes | NO | Newborn/term | NA |
|  | 7<sup>th</sup> | LRV/16 854 | + | NA | I | IgM-/IgG- | + | - | - | Third trimester / villous<br>immaturity | NO | Newborn/term | NA |
|  | 7<sup>th</sup> | LRV/16 859 | - | - | + | IgM-/IgG+ | + | - | - | Third trimester / without<br>pathological changes | NO | Newborn/term | NA |
|  | 7<sup>th</sup> | LRV/16 931 | + | NA | I | IgM-/IgG- | - | - | - | Third trimester / without pathological changes | NO | Newborn/term | NA |
|  | 7<sup>th</sup> | LRV/16 848 | NA | NA | NA | NA | + | - | - | Third trimester / without pathological changes | NO | Newborn/term | NA |
|  | 8<sup>th</sup> | LRV/16 284 | - | + | + | IgM-/IgG+ | - | NT | + | Third trimester / without pathological changes | NO | Newborn | Serum RT-PCR-/IgM- |
| UNKNOWN | ? | LRV/16 857 | NA | NA | NA | NA | + | - | - | Third trimester / without pathological changes | YES – Hydrocephalus | Newborn/preterm (34 w) | NA |
|  | ? | LRV/16 103 | NA | NA | NA | NA | + | - | - | Third trimester / villous immaturity | YES – Stillborn | Intrauterine fetal demise/term | Viscera RT-PCR- |
|  | ? | LRV/16 515 | NA | NA | NA | NA | + | - | - | Third trimester / villous immaturity | YES – brain abnormalities, encephalocele, death at 1-month-old | Newborn/term (38 w) | Viscera and CSF RT-PCR- |
|  | ? | HC N16-09 | NA | NA | NA | NA | + | - | NA | Third trimester / villous immaturity | YES – Microcephaly, discrete multifocal cerebritis, calcifications | Intrauterine fetal demise/term | NA |
|  | ? | LRV/16 855 | NA | - | NA | NA | + | - | - | Third trimester / without pathological changes | YES – Hydrocephalus, ventriculomegaly, calcifications, bilateral congenital cataract | Newborn/preterm (34 w) | CSF RT-PCR- |

\*For term placentas, the probable month of infection was calculated assuming delivery at week 39 as described by Moore et al. (Moore et al., 2014). †IHC was positive in the brain tissue of the newborn. ?, not known; +, positive; -, negative; I, inconclusive; NT, not tested; NA, not available; w, weeks; CSF, cerebrospinal fluid.

Serological analysis was performed in the 15 cases for which serum samples were available; nine of these cases presented with both anti-ZIKV IgM and anti-DENV IgG, and one had only anti-DENV IgG. The presence of these IgG antibodies suggests a past *Flavivirus* infection. None of the anti-ZIKV IgM-positive samples crossreacted in the anti-DENV IgM ELISA (Table 1, cases LRV/16 1318, 16 1317, 16 870, 16 1065, 16 1068, 16 1029, 16 1004, 16 859 and 16 284).

Neither ZIKV RNA nor anti-ZIKV IgM were detected in samples of amniotic fluid, newborn cerebrospinal fluid (CSF) or ocular/oral swabs in the cases described in this article. Of the five newborn serum samples available, one presented anti-ZIKV IgM, indicating transplacental infection, although no congenital disorder was observed at the time of delivery in this case (Table 1, case LRV/16 1065).

The anatomopathological findings for the FFPE placental tissue specimens are compiled in Tables 1 and 2. Of note, although our panel includes samples from women who had ZIKV infection at different stages of pregnancy, all placental samples were obtained at the time of delivery except for one, which was from a spontaneous abortion at 12 weeks of gestation (first trimester placenta, case LRV/15 572). Additional information about this case has been published (Noronha et al., 2016). This ZIKV-positive first trimester human placental sample (LRV/15 572) exhibited chronic villitis and TORCH-like features that are shown in Table 2 and Figure 1A.

**Figure 1.**
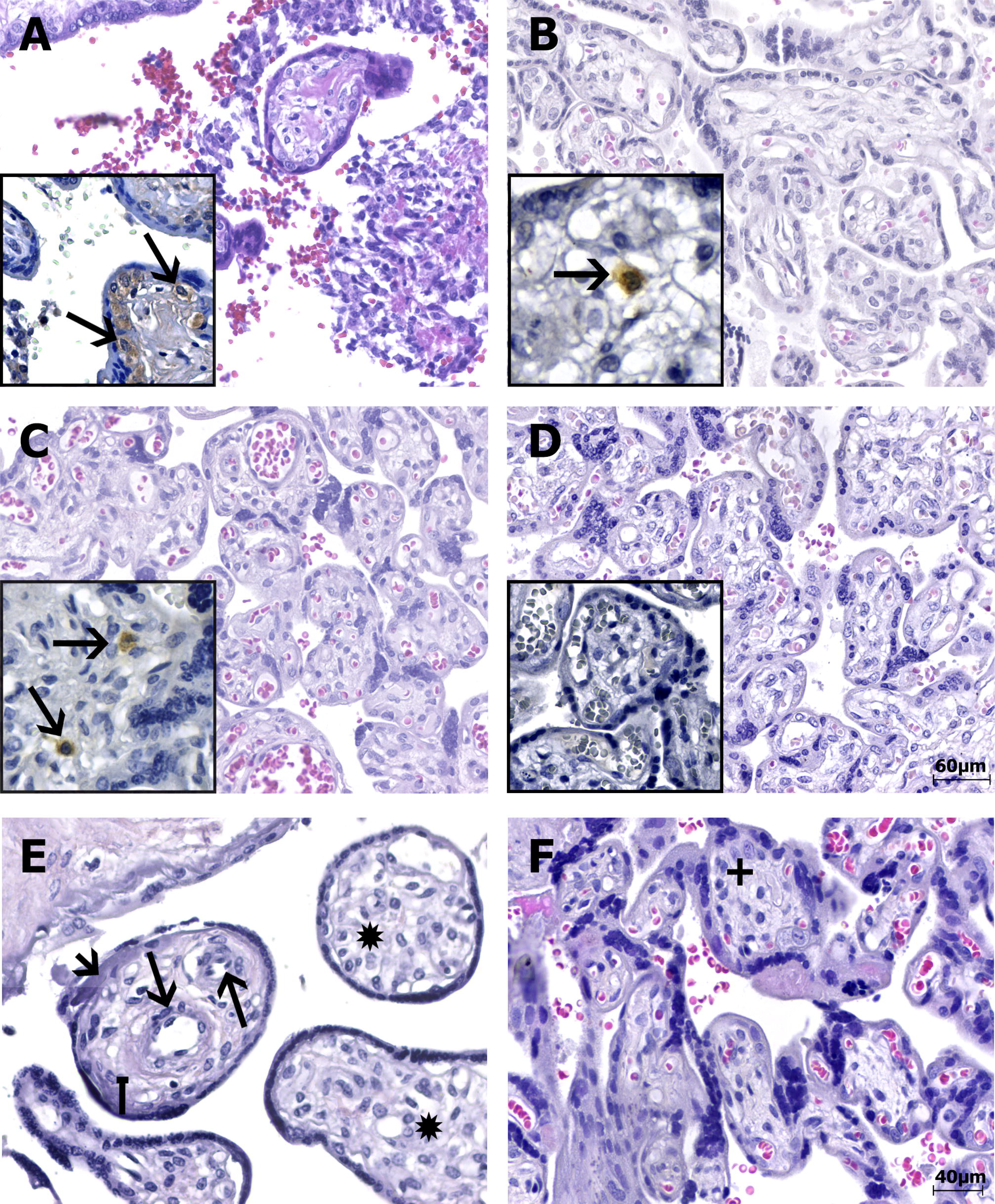
Photomicrography of the placental samples stained with H&E or immunostained with anti-ZIKV MAb and stained with Harris’s hematoxylin (squares). The scale bars are 60 μm **(A-D)** and 40 μm **(E-F)**. **(A)** Case LRV/15 572 – First trimester placenta sample (chorion frondosum) showing chronic villitis (TORCH-like) with lymphohistiocytic chronic villous inflammation. **(B)** Case LRV/16 515 – Third trimester placenta sample (chorion frondosum) showing delayed villous maturation with additional stromal changes such as stromal fibrosis. **(C)** Case LRV/16 845 – Third trimester placenta sample (chorion frondosum) showing delayed villous maturation with persistence of the cytotrophoblastic layer, stromal fibrosis and reduced numbers of syncytial knots. **(A-C)** The squares highlight Hofbauer cells positive for anti-ZIKV MAb (arrows) inside the chorionic villi. Notice that the cytotrophoblast and syncytiotrophoblast cells, as well as other fibroblastic cells inside Wharton’s jelly, are negative for anti-ZIKV MAb. **(D)** Negative control (ZIKV-negative patient) – Third trimester placenta sample (chorion frondosum) without pathological changes. Hofbauer cells are negative for anti-ZIKV MAb inside the chorionic villi (square). Note that the cytotrophoblast and syncytiotrophoblast cells, as well as other fibroblastic cells inside Wharton’s jelly, are negative for the antibody used. **(E)** Case LRV/15 578 – Third trimester placenta sample (chorion frondosum) showing delayed villous maturation with additional stromal changes, such as stromal fibrosis (square arrow), an increase in the number of fetal capillaries (arrows), hyperplasia of Hofbauer cells (*) and an increase in intravillous fibrinoid deposits (arrow head). **(F)** Negative control (ZIKV-negative patient) – Third trimester placenta sample (chorion frondosum) showing a normal number of Hofbauer cells (+). No delayed villous maturation or additional stromal changes were observed.

**Table 2.**
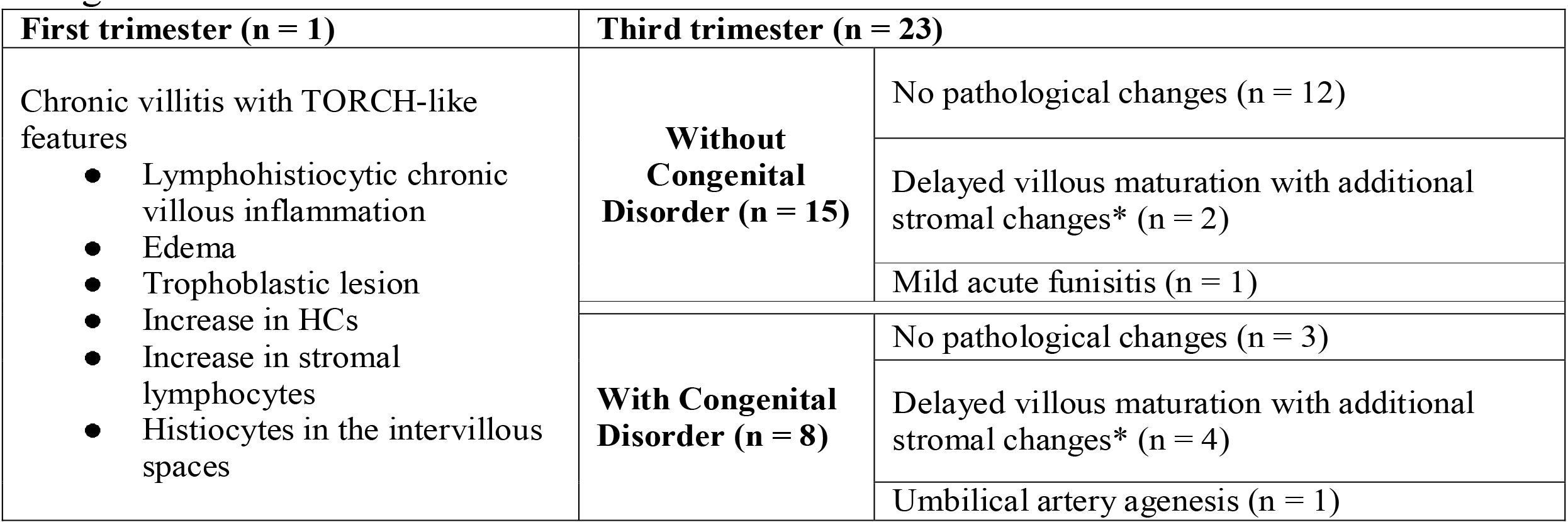
Main placental histopathological findings from infants with and without congenital disorder at birth.

Eight of the 23 third trimester human placental tissue samples exhibited pathological features in routine H&E-stained sections (whereas placentas without pathological signs were observed in the remaining 15 cases). The most prevalent alteration was delayed villous maturation with additional stromal changes, such as hyperplasia of HCs (75%), as shown in Table 2 and Figures 1B, 1C and 1E. Of the nine cases exhibiting pathological signs in the H&E staining, five were associated with congenital disorders (55.6%), and one evolved to a spontaneous abortion (first trimester placenta, case LRV/15 572). In the remaining 15 cases, no pathological evidence was observed in the H&E sections, but three of these cases presented with congenital disorders (20%) (Tables 1 and 2). Of the six cases with villous immaturity, four had a congenital disorder diagnosed at birth (66.7%), two had intrauterine fetal death (Table 1), one had acute funisitis, and one had umbilical artery agenesis (Table 2).

Morphometric analysis of the placentas from women infected during the third trimester of pregnancy confirmed the villous immaturity and hyperplasia of HCs (Figure 2). Syncytial knots and sprouts were quantified to demonstrate disorders of villus maturation, and a higher number of sprouts was detected in the ZIKV-infected group than in the control group (*p*<0.05) (Figure 2B). Villi and CD163+ HCs were measured to evaluate hyperplasia of HCs. Compared to the placentas in the control group, the placentas from women infected with ZIKV during the third trimester showed increased numbers of HCs (hyperplasia) (Figure 2C-D).

**Figure 2.**
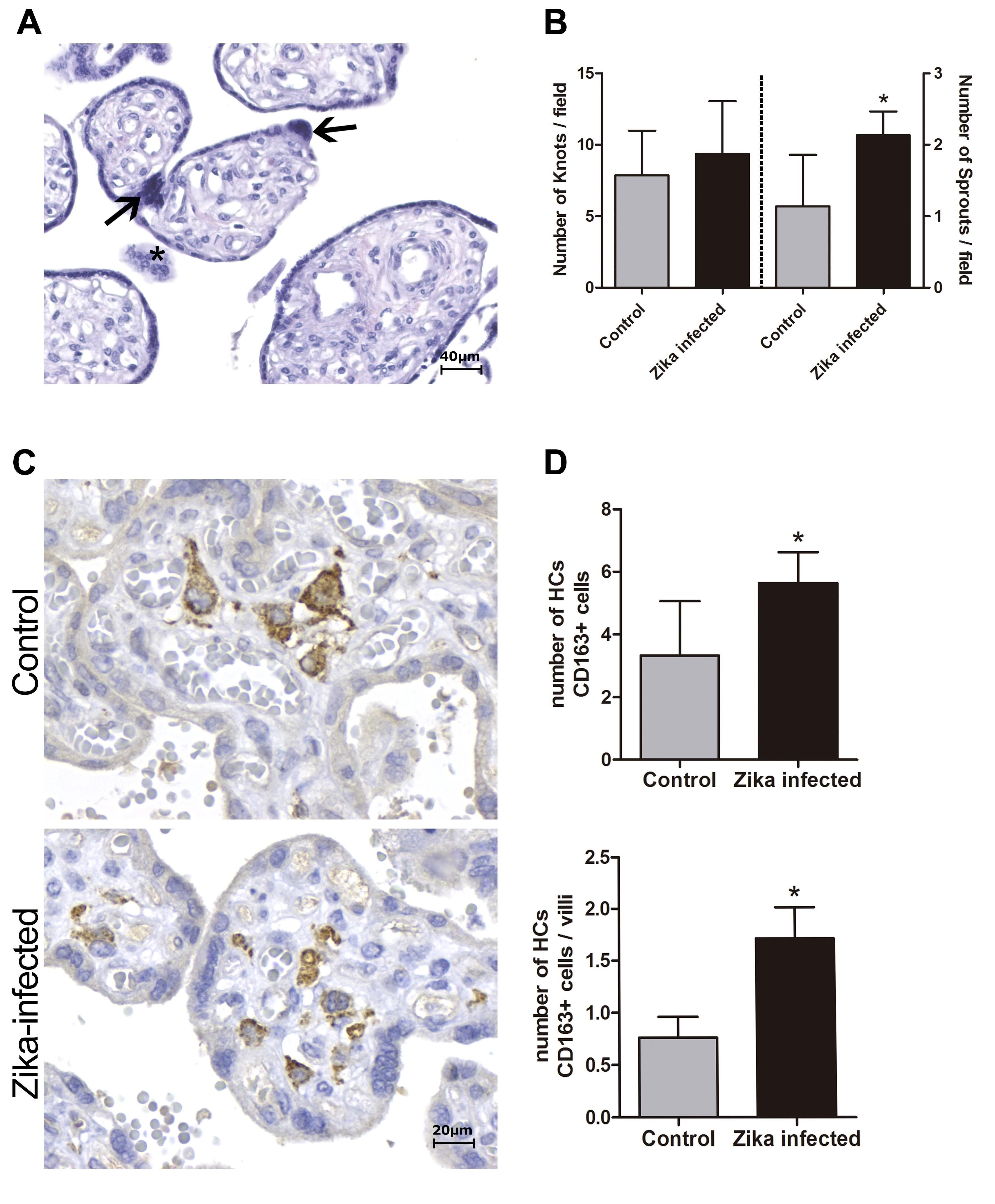
Morphometric analysis of placental specimens from women infected with ZIKV during the third trimester of pregnancy and from negative controls. **(A)** Photomicrography of a placental sample stained with H&E showing syncytial knots (arrows) and sprouts (*). **(B)** Quantification of knots and sprouts. A significant increase in the number of placental sprouts was observed in the ZIKV-positive group (n=6) compared to the negative controls (n=5). **(C)** Immunostaining with CD163 highlights Hofbauer cell hyperplasia in ZIKV-infected placentas. **(D)** Quantification of CD163+ Hofbauer cells. The average numbers of CD163+ cells and CD163+ cells per villus were significantly higher in the ZIKV group (n=5) than in the negative controls (n=3). **(B and D)** Thirty high-power fields (HPFs = 400×) for each case were randomly selected for counting. The mean of the 30 fields was used for the statistical analyses. The Zika-infected and negative control groups were compared using the Mann-Whitney test. The asterisks indicate statistically significant differences between the groups (*p*<0.05).

Immunohistochemical (IHC) analysis of the placental tissue samples using anti-flavivirus MAb (4G2) and anti-ZIKV MAb showed immunostaining in the HCs, regardless of the gestational age when ZIKV infection occurred (Figures 1A-C and 3E-F). The cytotrophoblast (CTB) and syncytiotrophoblast (STB) cells (chorion frondosum), as well as the fibroblastic cells and Wharton’s jelly, were negative in all specimens (Figures 1 and 3E-F). All umbilical cord (fibroblasts, Wharton’s jelly, and amniotic epithelium), chorioamniotic membrane (capsular decidua, amniotic epithelium, and smooth chorion above the capsular decidua), and decidua basalis samples were consistently negative for the anti-ZIKV and anti-flavivirus monoclonal antibodies (Figure 3A-C). The intervillous space showed maternal inflammatory cell infiltrates that were positive for both the anti-flavivirus and anti-ZIKV MAbs in only the first trimester placental sample (LRV/15 572) with villitis (Figure 3D). ZIKV infection of this sample was confirmed by RT-qPCR using RNA extracted from a FFPE tissue core corresponding to a villitis area (Noronha et al., 2016).

**Figure 3.**
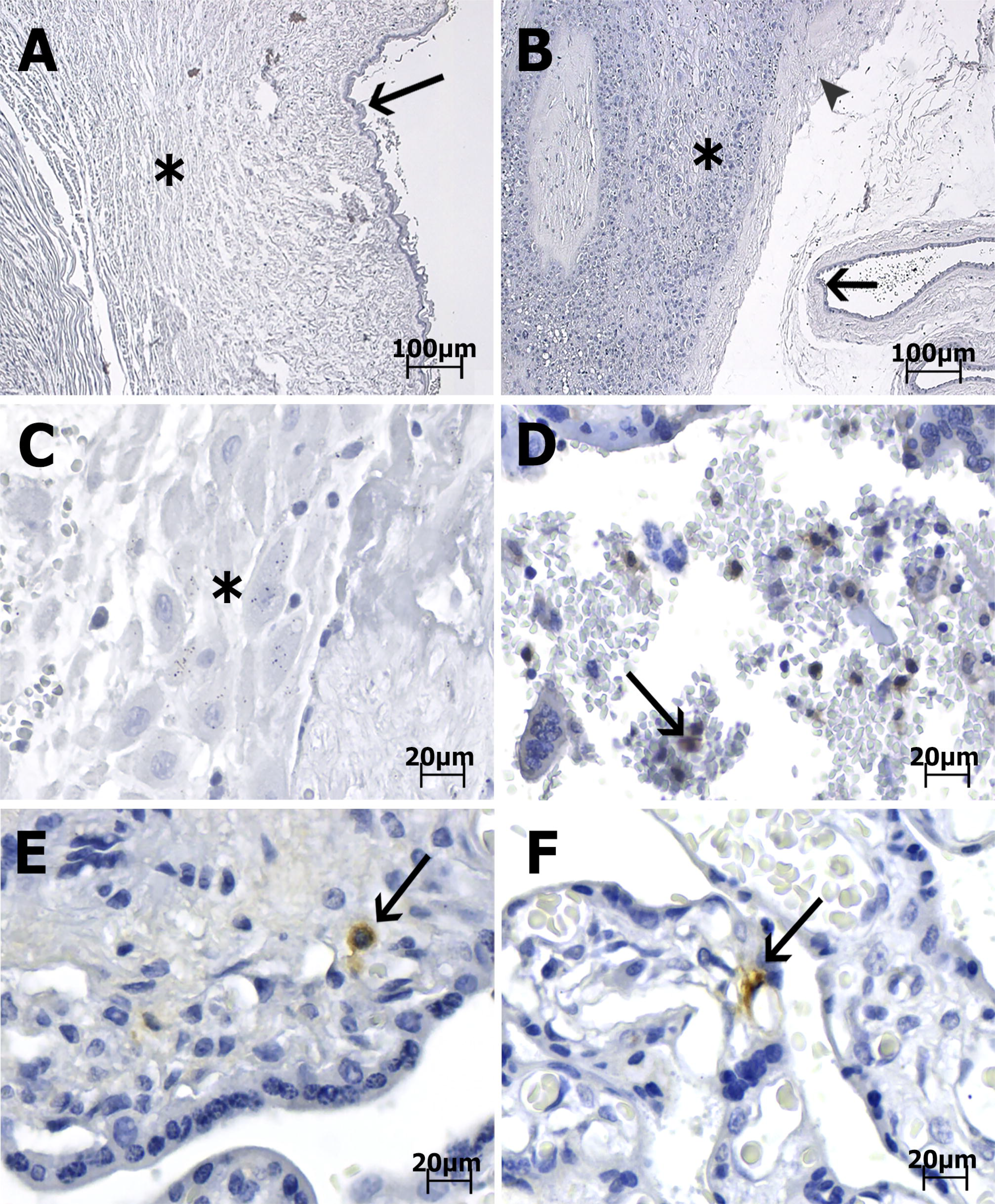
Photomicrography of placenta, cord and membrane samples from women diagnosed as positive for ZIKV infection immunostained with anti-ZIKV MAb and stained with Harris’s hematoxylin. The scale bars are 100 μm **(A-B)** and 20 μm **(C-F)**. **(A)** Case LRV/16 515 – Umbilical cord slides from a third trimester placenta negative for anti-ZIKV MAb. Fibroblasts in Wharton’s jelly (*) and amniotic epithelium (arrow) are negative for the antibody used. **(B)** Case LRV/16 515 – Chorioamniotic membrane from a third trimester placenta negative for anti-ZIKV MAb. Notice that the capsular decidua (*), the smooth chorion above the capsular decidua (arrowhead) and the amniotic epithelium (arrow) are negative for the antibody used. **(C)** Case LRV/16 515 – Decidua basalis (*) from a third trimester placenta negative for anti-ZIKV MAb. **(D)** Case LRV/15 572 – Intervillous space from a first trimester placenta with inflammatory cell infiltrates positive for anti-ZIKV MAb (arrow). **(E)** Case LRV/16 927 and **(F)** Case LRV/16 848: Chorion frondosum from third trimester placentas showing Hofbauer cells positive for anti-ZIKV MAb (arrows) inside the chorionic villi. Notice that the cytotrophoblast and syncytiotrophoblast cells, as well as fibroblastic cells inside Wharton’s jelly, are negative for the antibody used.

## 4 Discussion

In this study, we examined human placental samples from naturally infected women, focusing on the anatomopathological and morphological aspects of the samples and on determining which cells were targeted by ZIKV during pregnancy.

The main histological findings (from H&E sections) in the placental tissues from women that were infected with ZIKV at different gestational periods were villous immaturity and stromal changes, such as hyperplasia of HCs, stromal fibrosis, edema of stromal villi, sclerotic villi, calcification foci and fibrinoid deposits (Table 2). Although studies on naturally infected placental tissues are limited, two previous reports corroborate our findings (Martines et al., 2016b; Ritter et al., 2017). Both reports detected chorionic villi with calcification, fibrosis, perivillous fibrin deposition, patchy intervillositis and focal villitis in the cases that presented pathological signs. Conversely, tissues from several ZIKV-positive samples had normal-appearing chorionic villi, in agreement with the results shown in Table 2 (Martines et al., 2016b; Ritter et al., 2017).

Morphometric analysis showed enhancement of the number of HCs and syncytial sprouts (indicating a villous maturation disorder) in the placentas of women infected with ZIKV at late gestational periods compared to the numbers of these features in the control group placentas. Syncytial knots are specializations in the syncytiotrophoblast, and an increase in these structures in late gestation indicates placental pathology and can be used to evaluate villous maturity. Syncytial sprouts are markers of trophoblast proliferation; they are seen frequently during early pregnancy and are increased in many diseases (Loukeris et al., 2010). These findings agree with the results of a study showing the damaging effect of type I interferon (INF I) on *ex vivo* human mid-gestation placental tissues treated with INF-β. The authors demonstrated that type I INFs trigger fetal death in a mouse model and that treatment of *ex vivo* human placentas with INF-β induces morphological changes in the villi, such as syncytial knots and sprouts (Yockey et al., 2018).

Immunohistochemical analysis revealed that HCs were the only ZIKV-positive fetal cells in the naturally infected human placental samples examined in this study, regardless of the gestational period at which infection occurred, and that HCs remain persistently infected until the time of delivery. It is noteworthy that even in the tissue samples from women infected in late pregnancy, i.e., with a short interval between the acute phase of infection and delivery time (LRV/16854, 16859, 16931, 16848, 16927 and 16284), ZIKV was detected exclusively in HCs (Table 1).

HCs are placental villous macrophages of fetal origin, and alterations in their numbers (hyperplasia) and biological features are associated with complications in pregnancy. HCs play a role in diverse functions such as placental vasculogenesis, immune regulation and the secretion of enzymes and cytokines across the maternal-fetal barrier. In addition, the results from a double-labeling assay showed the association of Sprouty proteins (implicated in villus branch morphogenesis) with HCs, suggesting the involvement of HCs in the development of placental villi (Anteby et al., 2005).

HCs are categorized as placental M2 macrophages, and their location and migratory behavior confer an ability to move around the villous stroma and to make transient contacts with other macrophages and villous core cells (Khan et al., 2000; Tang et al., 2011).

Another fact worth mentioning is the apparent correlation between villous immaturity and congenital disorders caused by ZIKV infection since we observed the occurrence of fetal malformations in four out of six cases (66.7%) of villous immaturity (observed in H&E staining and morphometric analysis). Furthermore, villous immaturity may be related to an increase in HCs, the cells that sustain the presence of ZIKV in the placenta. These data could be valuable to pathologists and neonatologists following up on newborns exposed to ZIKV by vertical transmission.

Of note, the case of the first trimester placenta with villitis (LRV/15 572) exhibited ZIKV-positive maternal inflammatory cells in the intervillous space. Although positivity in fetal endothelium and maternal leukocytes (Ritter et al., 2017), and in decidual cells, CTBs and mesenchymal cells of chorionic villi (Rabelo et al., 2018) in a few first trimester placentas has also been reported, we speculate that maternal or other fetal cells may be infected only transiently. In support of this hypothesis, studies of natural ZIKV infections using either IHC or in situ hybridization analysis (ISH) have also reported that HCs were the most frequently observed infected cells (Bhatnagar et al., 2017; Martines et al., 2016a; Noronha et al., 2016; Rosenberg et al., 2017).

The persistence of ZIKV-positive fetal HCs in full-term placentas regardless of the period at which infection occurred indicates that ZIKV can persist in the placenta for several months after maternal infection and may provide a viral source for continued fetal infection.

Examination of the entire human placenta, comprising the umbilical cord, amniotic membrane, chorion frondosum (CTB and STB), smooth chorion, capsular decidua, and decidua basalis, revealed that all these tissues were consistently negative for ZIKV infection. Mlakar et al. (Mlakar et al., 2016) reported similar findings in a single case. Rabelo et al. (Rabelo et al., 2017) showed ZIKV NS1 protein in the decidual and endothelial cells of the maternal decidua and in CTB, STB, and HCs in the third trimester placental tissues associated with an HIV-exposed but uninfected infant with severe congenital Zika syndrome. Nonetheless, the maternal HIV infection could have contributed to the permissiveness of other maternal/placental cell types to ZIKV infection.

Thus, we suggest that the most plausible hypothesis for the transplacental transmission of ZIKV would be related to its association with HCs and its migratory ability to reach the fetal vessels and then infect the fetus either by transcytosis or through ZIKV-infected “Trojan horse” cells (Zanluca et al., 2018).

Furthermore, as both ZIKV genomic material and viral particles are detected in placental cells until the end of pregnancy (regardless of the trimester in which infection occurred), it is plausible to speculate that the infection of the fetus could happen as a secondary event, i.e., not necessarily concomitant with the maternal acute phase of disease (Aagaard et al., 2017). In this case, understanding the biology of HCs after ZIKV infection is of utmost importance in explaining the different congenital outcomes related to ZIKV infection (Simoni et al., 2017).

We emphasize that a negative ZIKV detection (by RT-PCR or IHC) in a placental sample does not exclude the possibility of maternal ZIKV infection. Possible reasons for false negative results include ZIKV RNA/protein levels below the limit of detection of the employed assays, RNA degradation due to storage/shipping processes or variability in tissue sampling, and the viral strain. For example, eight of the pregnant women described in Table 1 had ZIKV infection confirmed by RT-qPCR in the serum sample during the acute phase of infection, but no viral RNA/protein was detected in their placental tissues.

Conversely, in this study, ZIKV was also detected in placental samples from nine women who had an onset of ZIKV clinical symptoms during the first, second or third trimester but gave birth to normal infants. Abnormalities in these infants may have been prevented because placental integrity limited viral spreading from mother to fetus. However, we cannot exclude fetal exposure to ZIKV, and it is noteworthy that, in some cases, abnormalities are only detected months after delivery (Aragao et al., 2017; Ventura et al., 2017). Serological examination and clinical follow-up of the newborns are required to confirm and define the diagnosis. Periodic monitoring of these infants may be helpful for the early recognition of future sequelae from congenital infection (Bhatnagar et al., 2017). In conclusion, tissue analysis provides the opportunity to confirm maternal/placental ZIKV infection, which may alert the practitioner to the potential for a congenital disorder in the child.

## Acknowledgments

We thank Dr. Margareth MacDonald from Rockefeller University for the critical reading of the manuscript. We also thank Dr. Mirian Woiski, the Health Surveillance team and the "Mãe Paranaense" initiative from Secretaria de Estado da Saúde do Paraná. We thank Elizabeth Droppa, Célia Cruz, Carla S. Felippe and the LACEN-PR group for their support in clinical sample collection and shipping. We express our gratitude to Wagner Nagib for graphical design and to Associação Amigos do HC for logistic support.

Financial support: Fiocruz, CNPq, Capes Zika Fast Track and Fundação Araucária. CNDS and LN have a CNPq fellowship.

## Author contributions

LN, CZ, MB and CNDS contributed to the study design; CZ, AAS and MA performed the experiments; PZR performed the morphometric analyses; and MB, IMN and LST participated in the patient follow-up. MMP participated in sample identification/distribution. LN, CZ, AAS and CNDS analyzed the results and wrote the manuscript.

---

**Supplementary Figure 1.** Characterization of the anti-ZIKV MAb by immunofluorescence. The MAb recognized ZIKV-infected C6/36 cells and showed no crossreactivity with DENV serotypes 1 to 4 or with the yellow fever (YFV), West Nile (WNV) and Saint Louis encephalitis (SLEV) viruses. No reaction was observed in the MOCK-infected cells. The pan-flavivirus MAb 4G2 was used as the positive control, and an unrelated MAb was used as the negative control. The scale bars are 250 μm.

**Supplementary Figure 2.** Photomicrography of third trimester placental samples (chorion frondosum) from women diagnosed positive for ZIKV infection immunostained with anti-ZIKV **(A)**, anti-pan-flavivirus 4G2 **(B)** and anti-CHIKV MAbs **(C)** or with no primary antibody **(D)** and stained with Harris’s hematoxylin. **(A–B)**: The arrows indicate positive Hofbauer cells inside the chorionic villi. Notice that the cytotrophoblast and syncytiotrophoblast cells, as well as fibroblastic cells inside Wharton’s jelly, are negative for both antibodies used. **(C–D)**: No reaction was observed in the negative controls. The scale bars are 60 μm.

## References

Aagaard, K. M., Lahon, A., Suter, M. A., Arya, R. P., Seferovic, M. D., Vogt, M. B., et al. (2017). Primary human placental trophoblasts are permissive for Zika virus (ZIKV) replication. Sci. Rep. 7, 41389. doi:10.1038/srep41389.

Anteby, E. Y., Natanson-Yaron, S., Greenfield, C., Goldman-Wohl, D., Haimov-Kochman, R., Holzer, H., et al. (2005). Human placental Hofbauer cells express sprouty proteins: a possible modulating mechanism of villous branching. Placenta 26, 476–483. doi:10.1016/j.placenta.2004.08.008.

Aragao, M., Holanda, A., Brainer-Lima, A., Petribu, N., Castillo, M., van der Linden, V., et al. (2017). Nonmicrocephalic infants with congenital Zika syndrome suspected only after neuroimaging evaluation compared with those with microcephaly at birth and postnatally: How large is the Zika virus “iceberg”? AJNR Am J Neuroradiol 38, 1427–1434.

Baurakiades, E., Martins, A., Victor Moreschi, N., Souza, C., Abujamra, K., Saito, A., et al. (2011). Histomorphometric and immunohistochemical analysis of infectious agents, T-cell subpopulations and inflammatory adhesion molecules in placentas from HIV-seropositive pregnant women. Diagn Pathol 6, 101.

Beckham, J. D., Pastula, D. M., Massey, A., and Tyler, K. L. (2016). Zika virus as an emerging global pathogen. JAMA neurol 73, 875–879. doi:10.1001/jamaneurol.2016.0800.Zika.

Bhatnagar, J., Rabeneck, D. B., Martines, R. B., Reagan-Steiner, S., Ermias, Y., Estetter, L. B. C., et al. (2017). Zika virus RNA replication and persistence in brain and placental tissue. Emerg. Infect. Dis. 23, 405–414. doi:10.3201/eid2303.161499.

Brasil, P., Pereira, J. P., Moreira, M. E., Ribeiro Nogueira, R. M., Damasceno, L., Wakimoto, M., et al. (2016). Zika Virus Infection in Pregnant Women in Rio de Janeiro. N. Engl. J. Med. 375, 2321–2334. doi:10.1056/NEJMoa1602412.

CDC. Centers for Disease Control and Prevention (2017). Zika MAC-ELISA. Available at: http://www.cdc.gov/zika/pdfs/zika-mac-elisa-instructions-for-use.pdf [Accessed July 6, 2017].

Duffy, M. R., Chen, T.-H., Hancock, W. T., Powers, A. M., Kool, J. L., Lanciotti, R. S., et al. (2009). Zika virus outbreak on Yap Island, Federated States of Micronesia. N. Engl. J. Med. 360, 2536–2543.

Emery, S. L., Erdman, D. D., Bowen, M. D., Newton, B. R., Winchell, J. M., Meyer, R. F., et al. (2004). Real-time reverse transcription – polymerase chain reaction assay for SARS-associated Coronavirus. Emerg. Infect. Dis. 10, 311–316.

ICTV. International Comittee on Taxonomy of Viruses (2017). Virus Taxonomy: 2016 Release. Available at: https://talk.ictvonline.org/taxonomy/ [Accessed March 12, 2017].

Johnson, B. W., Russell, B. J., and Lanciotti, R. S. (2005). Serotype-specific detection of dengue viruses in a fourplex real-time reverse transcriptase PCR assay. J. Clin. Microbiol. 43, 4977–4983. doi:10.1128/JCM.43.10.4977.

Khan, S., Katabuchi, H., Araki, M., Nishimura, R., and Okamura, H. (2000). Human villous macrophage-conditioned media enhance human trophoblast growth and differentiation in vitro. Biol Reprod 62, 1075–1083.

Lanciotti, R. S., Kosoy, O. L., Laven, J. J., Velez, J. O., Lambert, A. J., Johnson, A. J., et al. (2008). Genetic and Serologic Properties of Zika Virus Associated with an Epidemic, Yap State, Micronesia, 2007. Emerg. Infect. Dis. 14, 1232–1239.

Loukeris, K., Sela, R., and Baergen, R. N. (2010). Syncytial knots as a reflection of placental maturity: reference values for 20 to 40 weeks’ gestational age. Pediatr. Dev. Pathol. 13, 305–309. doi:10.2350/09-08-0692-OA.1.

Martines, R. B., Bhatnagar, J., de Oliveira Ramos, A. M., Davi, H. P. F., Iglezias, S. D., Kanamura, C. T., et al. (2016a). Pathology of congenital Zika syndrome in Brazil: a case series. Lancet 388, 898–904. doi:10.1016/S0140-6736(16)30883-2.

Martines, R. B., Bhatnagar, J., Keating, M. K., Silva-Flannery, L., Muehlenbachs, A., Gary, J., et al. (2016b). Evidence of Zika virus infection in brain and placental tissues from two congenitally infected newborns and two fetal losses - Brazil, 2015. Morb. Mortal. Wkly. Rep. 65, 1–2.

Melo, A. S. de O., Aguiar, R. S., Amorim, M. M. R., Arruda, M. B., Melo, F. de O., Ribeiro, S. T. C., et al. (2016). Congenital Zika virus infection: Beyond neonatal microcephaly. JAMA Neurol. 73, 1407–1416. doi:10.1001/jamaneurol.2016.3720.

Miner, J. J., and Diamond, M. S. (2017). Zika virus pathogenesis and tissue tropism. Cell Host Microbe 21, 134–142. doi:10.1016/j.chom.2017.01.004.

Mlakar, J., Korva, M., Tul, N., Popović, M., Poljšak-Prijatelj, M., Mraz, J., et al. (2016). Zika virus associated with microcephaly. N. Engl. J. Med. 374, 951–958. doi:160210140106006. Epub ahead of print.

Moore, K., Persuad, T., and Torchia, M. (2014). “Introduction to the developing human,” in The developing human: Clinically oriented embriology, eds. K. Moore, T. Persuad, and M. Torchia (Philadelphia: Elsevier), 48.

Noronha, L. De, Zanluca, C., Azevedo, M. L. V., Luz, K. G., and Duarte dos Santos, C. N. (2016). Zika virus damages the human placental barrier and presents marked fetal neurotropism. Mem Inst Oswaldo Cruz 111, 287–293. doi:10.1590/0074-02760160085.

Rabelo, K., de Souza Campos Fernandes, R. C., Souza, L. J. de, Louvain de Souza, T., Santos, F. B. dos, Guerra Nunes, P. C., et al. (2017). Placental histopathology and clinical presentation of severe congenital Zika syndrome in a human immunodeficiency virus-exposed uninfected infant. Front. Immunol. 8, 1–8. doi:10.3389/fimmu.2017.01704.

Rabelo, K., Souza, L. J., Salomão, N. G., Oliveira, E. R. A., Sentinelli, L. D. P., Lacerda, M. S., et al. (2018). Placental inflammation and fetal injury in a rare Zika case associated with Guillain-Barré Syndrome and abortion investigation of the placental tissue. Front. Microbiol. 9, 1018. doi:10.3389/fmicb.2018.01018.

Ritter, J. M., Martines, R. B., and Zaki, S. R. (2017). Zika virus: Pathology from the pandemic. Arch. Pathol. Lab. Med. 141, 49–59. doi:10.5858/arpa.2016-0397-SA.

Rosenberg, A. Z., Weiying, Y., Hill, D. A., Reyes, C. A., and Schwartz, D. A. (2017). Placental pathology of zika virus: Viral infection of the placenta induces villous stromal macrophage (Hofbauer Cell) proliferation and hyperplasia. Arch. Pathol. Lab. Med. 141, 43–48. doi:10.5858/arpa.2016-0401-OA.

Schuler-Faccini, L., Ribeiro, E. M., Feitosa, I. M. L., Horovitz, D. D. G., Cavalcanti, D. P., Pessoa, A., et al. (2016). Possible association between Zika Virus infection and microcephaly - Brazil, 2015. Morb. Mortal. Wkly. Rep. 65, 59–62.

Simoni, M. K., Jurado, K. A., Abrahams, V. M., Fikrig, E., and Guller, S. (2017). Zika virus infection of Hofbauer cells. Am. J. Reprod. Immunol. 77, e12613. doi:10.1111/aji.12613.

Tang, Z., Tedesse, S., Norwitz, E., Mor, G., Abrahamns, V. M., and Guller, S. (2011). Isolation of Hofbauer cells from human term placentas with high yield and purity. Am J Reprod Immunol 66, 336–348. doi:10.1111/j.1600-0897.2011.01006.x.Isolation.

Ventura, L., Ventura, C., Lawrence, L., van der Linden, V., van der Linden, A., Gois, A., et al. (2017). Visual impairment in children with congenital Zika syndrome. J AAPOS 21, 295–299.

WHO. World Health Organization (2017). Zika situation report. 2017. Available at: http://apps.who.int/iris/bitstream/10665/254714/1/zikasitrep10Mar17-eng.pdf?ua=1 [Accessed January 5, 2018].

Yockey, L. J., Jurado, K. A., Arora, N., Millet, A., Rakib, T., Milano, K. M., et al. (2018). Type I interferons instigate fetal demise after Zika virus infection. Sci. Immunol. 3, eaao1680. doi:10.1126/sciimmunol.aao1680.

Zanluca, C., de Noronha, L., and Duarte dos Santos, C. N. (2018). Maternal-fetal transmission of the zika virus: An intriguing interplay. Tissue Barriers 6, e1402143. doi:10.1080/21688370.2017.1402143.

Zanluca, C., and Duarte dos Santos, C. N. (2016). Zika virus – an overview. Microbes Infect. 18, 295–301. doi:10.1016/j.micinf.2016.03.003.

Zanluca, C., Melo, V. C. A. de, Mosimann, A. L. P., Santos, G. I. V. dos, Santos, C. N. D. dos, and Luz, K. (2015). First report of autochthonous transmission of Zika virus in Brazil. Mem. Inst. Oswaldo Cruz 110, 569–572.

